# A Practical Approach to Constructing a Knowledge Graph for Soil Ecological Research

**DOI:** 10.1101/2023.03.02.530763

**Authors:** Nicolas Le Guillarme, Wilfried Thuiller

## Abstract

With the rapid accumulation of biodiversity data, data integration has emerged as a hot topic in soil ecology. Data integration has indeed the potential to advance our knowledge of global patterns in soil biodiversity by facilitating large-scale meta-analytical studies of soil ecosystems. However, ecologists are still poorly equipped when it comes to integrating disparate datasets into a unified knowledge graph with well-defined semantics. This paper presents a practical approach to constructing a knowledge graph from heterogeneous and distributed (semi-)structured data sources. To illustrate our approach, we integrate several datasets on the trophic ecology of soil organisms into a trophic knowledge graph and show how information can be retrieved from the graph to support multi-trophic studies.

## Introduction

In recent years, a number of initiatives aiming at collecting new soil biodiversity data or assembling existing datasets have emerged, resulting in a rapid accumulation of data in soil ecology [1]. Because of the enormous phylogenetic, taxonomic and functional diversity of soil organisms, datasets are often collected by individual scientists or small project teams from different communities or disciplines to answer precise research questions. These datasets are typically small, with a limited spatial/temporal/taxonomic coverage, and are formatted according to the project needs, with little or no concern for data standardization [2]. This causes datasets to be heterogeneous in syntax (differences in models or languages), schema (differences in data structures and formats), and semantics (differences in terminologies, meaning or interpretation of data in different disciplines or research contexts). In addition, datasets are widely distributed: they reside on diverse locations, e.g. files or databases on the local network or published on the web, and are accessible using different interfaces, e.g., files download, database queries or web APIs.

Integrating these “long-tail data” dispersed across different datasets could help address research questions at larger scales [3]. Data integration is of growing interest in the ecological domain, with much efforts directed towards the creation of standard terminologies for describing, sharing and facilitating the aggregation of biodiversity data, e.g. organismal trait data [4, 5, 6, 7], into large open databases. Recent initiatives in trait-based ecology have targeted specific taxonomic groups, e.g. ants [8], spiders [9], soil invertebrates [10, 11], fungi [12, 13], plants [14]. Although these databases have made aggregated data more readily accessible to scientists, they are not yet interoperable. The difficulty of integrating data distributed across heterogeneous sources remains. As a result, integrative analyses of soil communities that span several taxonomic groups and integrate multitrophic interactions are scarce — see [15] for an example — although essential to improve our understanding of the links between soil biodiversity and ecosystem functioning [16].

Here, we address the problem of semantic data integration in the biodiversity science domain. Semantic data integration is the process of combining data from different sources into a single, unified view [17] (Figure 1). This provides the user with the ability to seamlessly manipulate data from multiple sources, regardless of the original format or location of the data. This “single, unified view” often takes the form of a knowledge graph.

**Figure 1.**
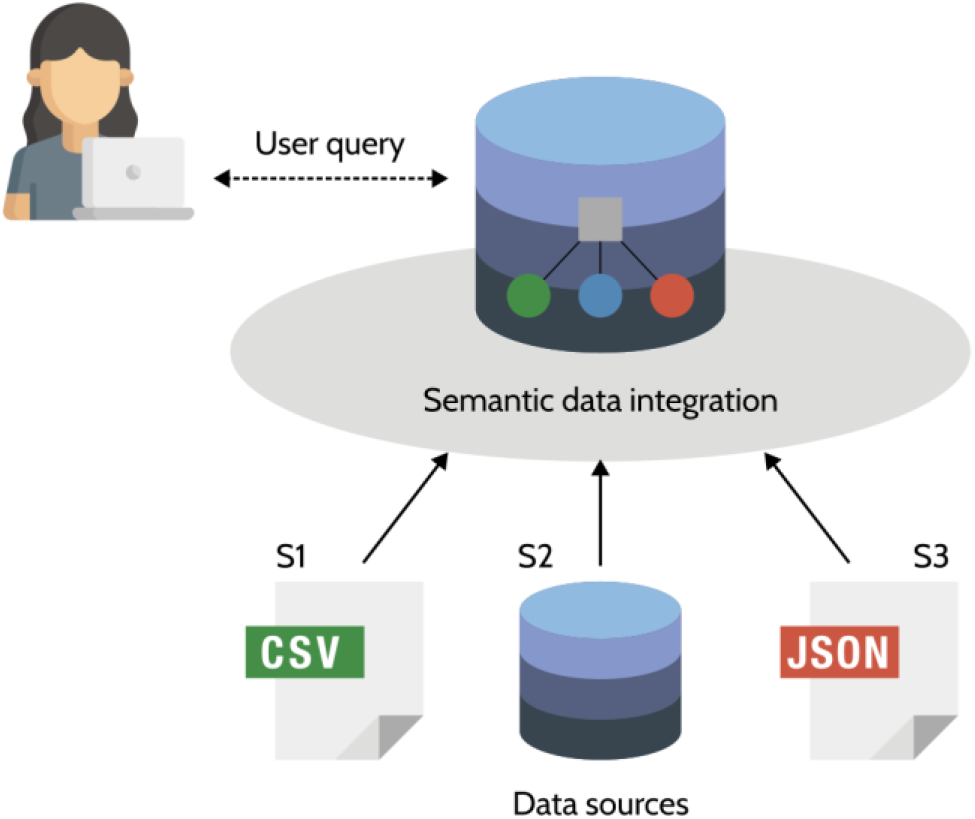
Semantic data integration provides the user with a uniform access to a set of autonomous and possibly heterogeneous data sources in a particular application domain.

A knowledge graph (KG) is a graph-structured knowledge base that stores factual information in the form of relationships between entities [18]. Under the Resource Description Framework (RDF), the standard data model of the Semantic Web, a KG is a set of (subject, predicate, object) triples. A RDF triple is a factual statement about an entity (the subject), connected to another entity or a data value (the object) by a relationship (the predicate). A set of RDF triples forms a labeled directed graph, called a RDF graph. But not every RDF graph is a KG. The triples in a KG can be separated into two distinct, yet connected, layers (Figure 2). The schema layer is the conceptual model of the KG and is described by an ontology or a collection of ontologies. An ontology is a formal conceptualization of a domain of interest [19]. It defines the types of entities that exist in the domain, their properties and relationships, using a logic-based ontology language — most often the Web Ontology Language (OWL), built upon RDF — that allows both humans and computers to understand the semantics of the data. The data layer holds the concrete, factual data. These data are instances of the general concepts defined in the ontology. In the context of semantic data integration, the data layer is populated with data from multiple sources. The ontology is used to link these disparate datasets at the schema-level, acting as a mediator for reconciling the structural and semantic heterogeneities between data sources.

**Figure 2.**
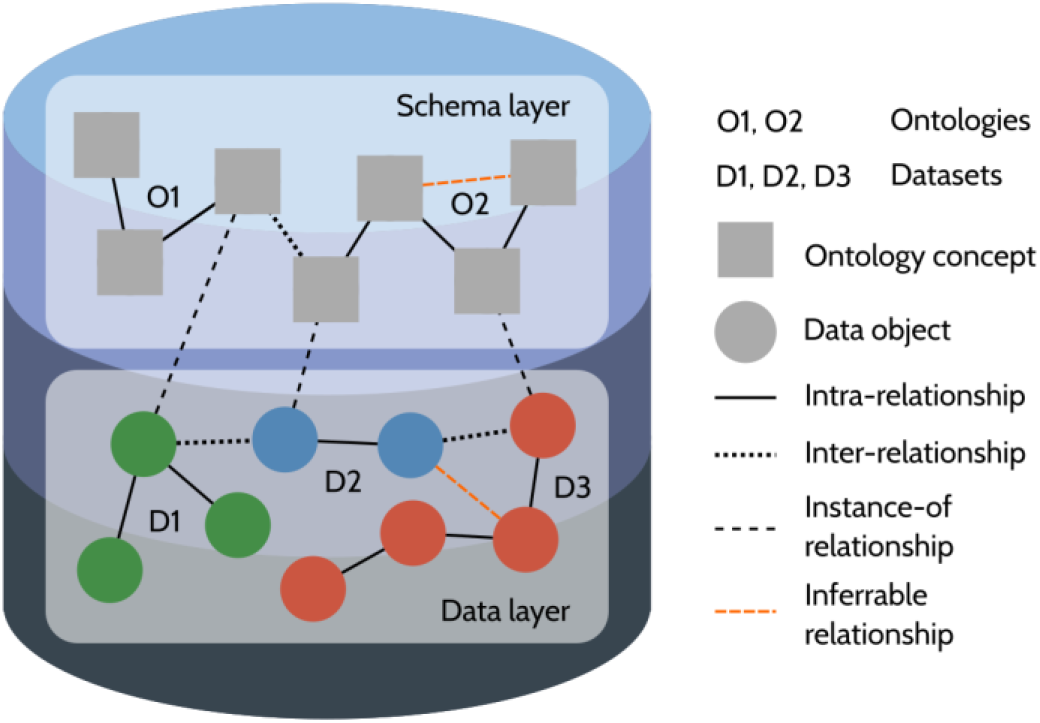
A knowledge graph is a graph database that embeds both the data and its semantics in two interconnected layers. The schema layer is an ontology or a collection of ontologies that integrate datasets at the schema-level and allow logical inference of implicit knowledge using a reasoner. The data layer is a collection of data from various sources.

KGs have a number of advantages over other types of databases, such as relational ones. Their graph structure allows for efficient querying, intuitive visualization, and analysis using graph algorithms or relational machine learning [18]. Using an ontology as a schema layer, KGs embed a formal semantics with the data which can be used by computers to interpret and reason about the data, thus potentially allowing to infer new facts (e.g. the inferrable relationships in Figure 2). KGs make it easy to integrate new types of data by altering the ontology or adding a new ontology to the schema layer. When following the Linked Open Data principles [20], domain-specific KGs can be easily interconnected into larger cross-domain KGs.

KGs have recently become prevalent as a framework for semantic data integration in many different domains of science and industry [21]. So far, only few examples of biodiversity KGs have been published. Ozymandias [22] is a KG for the Australian fauna that integrates data from several sources, including the Atlas of Living Australia, the Australian Faunal Directory, the Biodiversity Heritage Library and the Biodiversity Literature Repository. OpenBiodiv [23] integrates information extracted from the biodiversity literature into a graph database using the OpenBiodiv-O ontology and an RDF version of the Global Biodiversity Information Facility (GBIF) taxonomic backbone. TAXREF-LD [24] is a KG representation of the French national taxonomical register for fauna, flora and fungus that interlinks information about taxonomy, species interactions, development stages, biogeography, conservation statuses, etc. In a recent talk at TDWG 2021, Michel et al. [25] called for more biodiversity data producers to start publishing KGs. However, for now, building a KG from multiple data sources is a complex and time-consuming task that demands high Semantic Web expertise, and we are not aware of an existing tool specifically designed to help ecologists transform their data sets into interoperable KGs — with the notable exception of the iKNOW projet [26], which is very similar in spirit to our work, but whose current status is unknown to us.

In this paper, we present inteGraph, a framework and toolbox that facilitates the process of building a KG from heterogeneous and distributed (semi-)structured data sources in the biodiversity domain. InteGraph implements a declarative approach for creating automatic and reproducible semantic data integration pipelines, while minimizing the amount of manual efforts and Semantic Web expertise needed to turn datasets into interoperable KGs. To illustrate our approach, we will show how inteGraph can be used to integrate data on the trophic ecology of soil organisms from multiple sources into a KG that can support multi-trophic studies.

## Material and Methods

### Motivating example

Multitrophic studies, spanning multiple trophic levels and/or taxonomic groups, are essential to identify general patterns in community ecology [27], understand how diversity is related to ecosystem stability and ecosystem functioning [28], and provide the necessary guidance with biodiversity loss and environmental problems [29]. Multitrophic approaches should acknowledge the complexity of ecosystems while remaining practical. Large trait databases have the potential to address this trade-off between feasibility and completeness. By supporting the assignment of species (or higher taxonomic ranks) to trophic and/or functional groups, they reduce the dimensionality of ecological communities without biasing studies toward a single trophic level or taxonomic group [30, 15]. Yet, challenges remain: although we have trait databases available for some of them, our trait knowledge is limited for most groups of soil organisms. In addition, existing databases tend to function as data silos whose lack of interoperability can discourage researchers to include more trophic levels and/or taxonomic groups in their studies. Throughout this paper, we will show how our framework can be used to build a knowledge graph integrating trophic information from a number of trait databases covering different soil taxonomic groups across several trophic levels, thus providing a unified access to multigroup, multitrophic information.

### Overview of the approach

Figure 3 is a high-level representation of our declarative approach to constructing a KG from heterogeneous and distributed (semi-)structured data sources. The core of our framework is inteGraph^1^, a toolkit for ontology-based data integration in the biodiversity domain that allows generating data integration pipelines dynamically from configuration files and scheduling and monitoring the execution of these pipelines.

**Figure 3.**
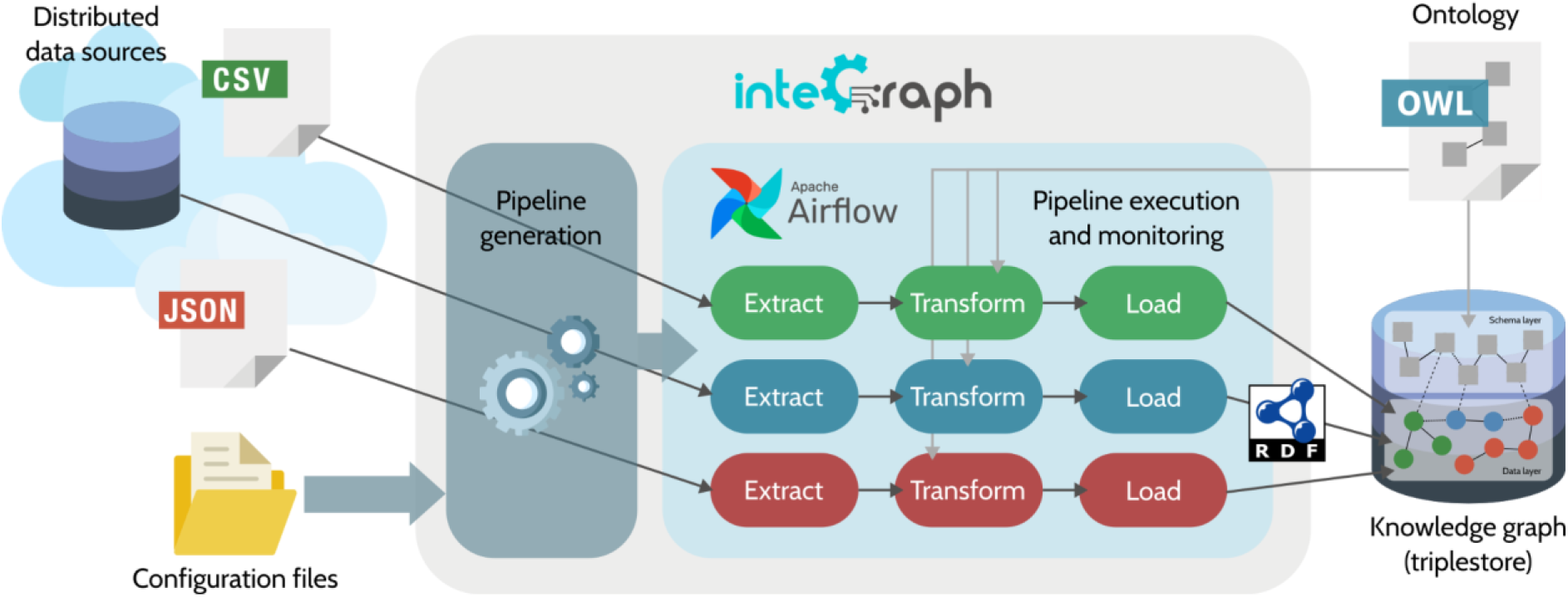
A high-level representation of the proposed declarative approach for constructing a knowledge graph from distributed (semi-)structured data sources.

### Data sources

Data sources can be (and often are) distributed on several machines, on the local network and/or on the web. Data must be accessible in a (semi-)structured form, for instance as tabular (e.g. tables in relational databases or in CSV files) or hierarchical data (e.g. data in XML or JSON format).

In our running example, we will use inteGraph to build a trophic KG from the following three data sources:

- The Fun^Fun^ database [13] collates fungal functional trait data, including information about trophic guilds, from a variety of sources, for thousands of species across the fungal tree of life. Data are provided in a tabular format (CSV file) and can be downloaded from Zenodo (https://zenodo.org/record/1216257).
- BETSI [11] is an open database gathering data on morphological traits and ecological preferences for 7 taxonomic groups of soil invertebrates (Aranae, Carabidae, Chilopoda, Collembola, Diplopoda, Isopoda and Diplotesticulata) from about 2000 literature references. BETSI is accessible on demand via a web portal (https://portail.betsi.cnrs.fr) that provides the user with an interface to write queries and download subsets of the database in a tabular format (CSV file). In the following, we will integrate a dataset containing Carabidae diet data.
- The Global Biotic Interactions (GloBI) provides open access to species interaction data (e.g. predator-prey, pollinator-plant, pathogen-host, parasite-host) aggregated from existing open datasets [31]. As of January 2023, GloBI contains over 15.5k interaction records obtained from 331 datasets, covering 888,544 taxa. GloBI provides several ways to access its data, including a web portal (https://www.globalbioticinteractions.org/), a downloadable snapshot of the entire database in a tabular format (CSV file), and a web API. In our example, we will use the web API to download data about the trophic interactions of Collembola.

### Target ontology

InteGraph adopts a top-down approach to KG creation. In this type of approach, the schema of the KG, i.e. the ontology, is defined upfront. The semantic data integration process then consists of populating the data layer of the KG by mapping the input data to the target ontology using mapping rules.

To reconcile schematic and semantic heterogeneities between our trophic data sources, we will use a global schema consisting of two ontologies: the NCBITaxon ontology and the Soil Food Web Ontology (SFWO) [7]. NCBITaxon is a formal translation of the NCBI Taxonomy database into an ontology, in which each taxon is treated as a class whose instances would be individual organisms, e.g. ‘Nicolas Le Guillarme’ instance_of NCBITaxon_9606 (*Homo sapiens*). Currently, the NCBI Taxonomy database is the only taxonomic nomenclature available as an ontology. SFWO is an ontology for representing knowledge on the trophic ecology of soil organisms across taxonomic groups and trophic levels. SFWO captures the semantics of trophic concepts such as trophic interactions, feeding processes, diets or trophic groups. SFWO also includes machine-interpretable definitions for most of these concepts, that allow for inference of implicit knowledge using automated reasoning, e.g. deducing a consumer’s diet(s) from the trophic interaction(s) in which it participates.

### Triplestore

A triplestore is a database management system, i.e., a software used for storing and querying a database, specifically designed to support the storage and the efficient querying of RDF data. A triplestore is needed to store both the schema and data layers that constitute a KG. Information stored in the triplestore can be retrieved using SPARQL queries. A multitude of triplestore implementations are available (see [32] for a survey), which offer different capabilities and performance in terms of data storage and indexing, query processing, reasoning, etc. InteGraph is not tied to a specific implementation and provides connectors to several triplestore solutions. As a top-down approach, inteGraph assumes that the target ontology has been loaded in the triplestore before starting the data integration process.

### Configuration files

As a declarative approach to building KGs, inteGraph provides control over the creation and execution of semantic data integration pipelines using configuration files. For each data source, the user must provide the following information:

- the type and the location of the data source;
- the credentials for connecting to the data source, if required;
- the format of the input data, e.g. tab or comma-separated values;
- the number of parallel processes;
- the structure of the input data, e.g. which columns contain taxonomic information;
- the schema mapping rules.

InteGraph also asks for global configuration files, including information to establish a connection with the triplestore, or to download the target ontology.

### Anatomy of inteGraph pipelines

InteGraph pipelines are structured according to the Extract-Transform-Load (ETL) paradigm. An ETL pipeline collects data from an input source (extract), cleans and maps the data from a source schema — the schema of the original data source — to a target schema (transform), and saves the transformed data into a target repository (load). In a typical ETL process, a copy of the extracted data is stored in a data staging area and all transformations are applied to the staged data. In inteGraph, an ETL pipeline is dynamically created at runtime for each data source from the configuration files provided by the user. This ETL pipeline extracts and stages the raw data from the data source, transforms the staged data into a RDF graph, and loads the RDF graph into the data layer of the triplestore.

### Data extraction

This first step of the ETL data integration process involves collecting data from the data source. InteGraph implements a number of components to connect to different types of data sources. At the moment, inteGraph supports the following types of data sources:

- File-like data sources: inteGraph can download files from remote or local file-like sources by specifying the URL of the source (starting with http://, https://, or file://) in the configuration. Archive files, including compressed archives, are supported, and unpacked before staging.
- HTTP data sources: inteGraph can extract data from remote databases exposed through a web-based API by sending HTTP GET requests to the API endpoint. In that case, the user is expected to provide the URL of the endpoint and the query string. Paginated results are supported using the limit and offset parameters.

These two components alone are sufficient to access most ecological datasets. We plan to add more connectors in the future, including connectors to SQL databases, RDF databases, etc. The extracted data are staged on the local file system in a tabular format.

Figure 4a shows the data extracted from our three example data sources. The three datasets use different data structures and terminologies to organize and describe taxonomic and trophic information.

- The Fun^Fun^ database uses the Index Fungorum taxonomic nomenclature. Each line of the data table contains a single trait information for a given taxon. The name of the trait is given in the trait_name column and its value(s) is given in the value column. The terminology used to describe the guild of each taxon is inherited from the FunGuild database.
- BETSI does not encode taxonomic information using identifiers from a reference taxonomy. Taxa are designated only by their scientific name. Similar to Fun^Fun^, each line of the data table contains a single trait information for a given taxon. The diet terminology is taken from the T-SITA thesaurus [4].
- Each line in GloBI’s data table contains information about a single interaction. The interaction is directed, so each line contains information about the source and target taxa (names and identifiers in a reference taxonomy) and the interaction name. GloBi maintains a mapping between different taxonomic nomenclature internally, but each taxon in the data table is linked to a single identifier. The target of the trophic interaction can also be a non-taxonomic entity, e.g. rotten wood.

### Data transformation

The second step of the ETL data integration process involves transforming the staged data into a RDF graph, i.e. a set of RDF triples. In inteGraph, data transformation consists of two successive operations: data cleansing and schema mapping.

Under the term data cleansing, we include all the dataset-specific data processing operations that aim at preparing the extracted data so that they are ready for further processing by the schema mapping component. This includes operations such as removing or filling missing values, removing duplicates, dropping irrelevant data, splitting strings (i.e. splitting a string representing a set of values, e.g. “bacterivore-detritivore”, into a set of strings, each string representing a single value, e.g. “bacterivore”, “detritivore”), joining two or more data tables, etc. Figure 4b shows an example of applying cleansing operations to the Fun^Fun^ dataset so that each line of the data table contains a single guild value. As possible cleansing operations are very diverse and highly dependent on the structure of the input data, inteGraph requires user intervention in the form of a Python or R script that specifies the operations that should be applied to the staged data before moving on to schema mapping. This script can be provided with the other source configuration files and is automatically incorporated in the ETL pipeline when created.

After data cleansing is complete (Figure 4b), cleansed data are staged and the pipeline moves on to schema mapping which involves converting data from the schema of the original data source to the schema of the knowledge graph, i.e. the target ontology. Schema mapping in inteGraph consists of two successive tasks: semantic annotation and RDF graph generation. Semantic annotation is the process of matching the input data with the concepts in the target ontology that best capture the semantics of the data. InteGraph provides two components for semantic annotation of biodiversity data (Figure 4c). The first component maps taxonomic data (taxon name and/or identifier in a source taxonomic nomenclature) to a target taxonomy — in our running example, the NCBITaxon ontology. Taxonomic mapping uses gnparser [33] to parse scientific names and nomer [34] to match taxon names and identifiers to their equivalent in the target taxonomy. The second component links non-taxonomic data, such as trait names or values, to concepts from the target ontology, e.g. linking the guild “Plant pathogen” to the concept “plant pathogen” whose identifier in the Soil Food Web Ontology is SFWO:0000159. Ontological mapping uses a naive approach based on text literals. The component retrieves all the concepts whose label (or the label of one of its synonyms) matches exactly the lookup term. If a single eligible concept is found, the term in the input data is annotated with this concept identifier. The user can also provide a dictionary containing term:concept pairs to handle mismatched terms or multiple matches. Figure 4c shows the result of applying semantic annotation on our example datasets.

**Figure 4.**
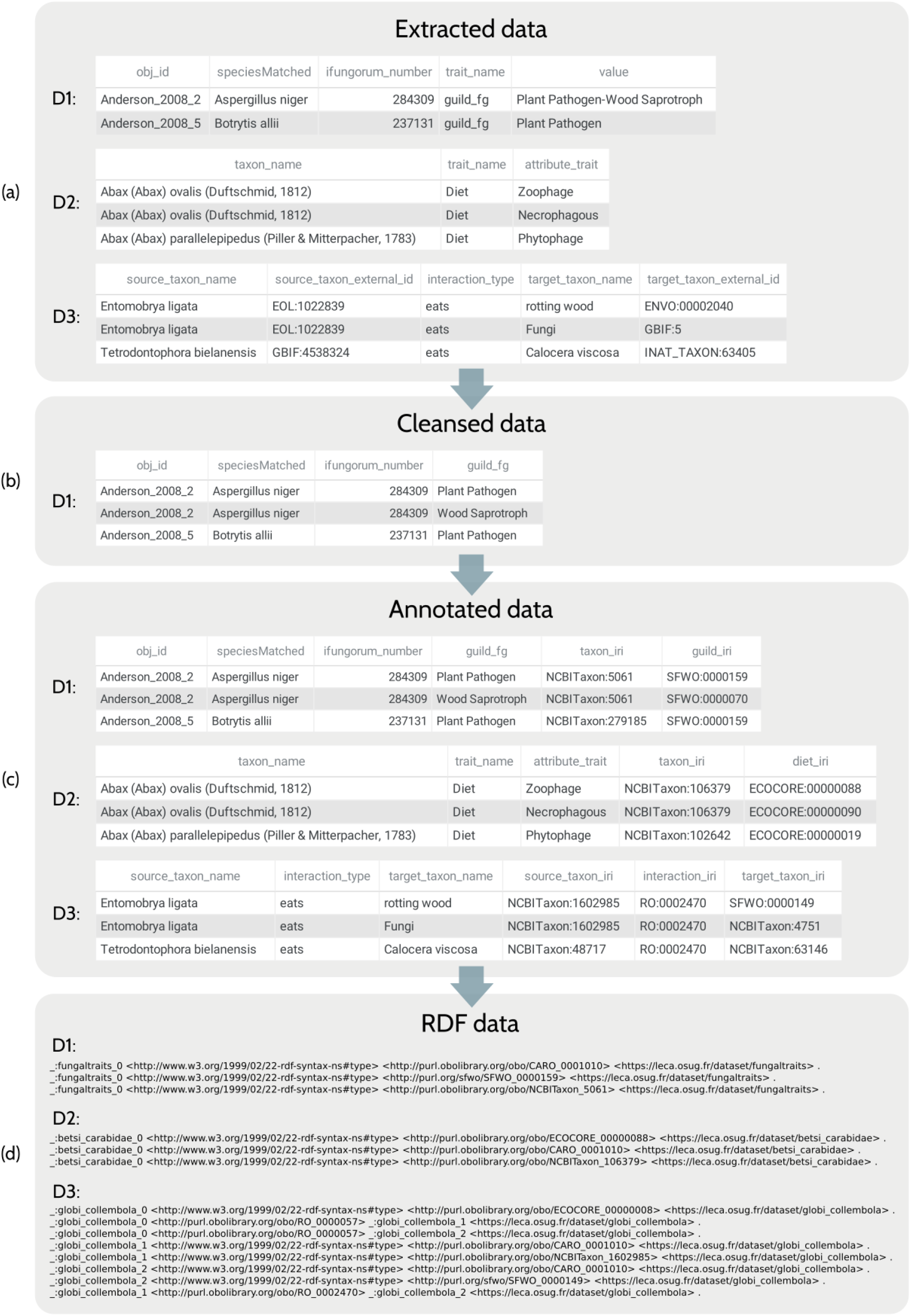
An illustration of the application of data transformation to our example datasets (D1: Fun^Fun^, D2: Carabidae diet data from BETSI, D3: Collembola trophic interaction data from GloBI). Figure 4d shows the set of RDF triples (in N-quads format) generated from the first line of each data table.

Once the relevant data have been linked to the corresponding concepts in the target ontology, the final step of schema mapping is the conversion of the annotated dataset into an RDF graph (Figure 4d). InteGraph uses RDF Mapping Language (RML) [35] rules to transform tabular data into RDF triples. RML is a declarative language for expressing rules that map data in heterogeneous structures to the RDF data model, the rules describing the desired graph structure. The schema mapping rules should be provided by the user as part of the data source configuration. However, writing RML mapping documents is beyond the reach of most non-expert users. To face this issue, inteGraph enables to specify mapping rules in spreadsheets that are automatically translated into RML documents using Mapeathor [36]. This provides a user-friendly manner to declare mapping rules in a language-independent way. Finally, inteGraph applies Morph-KGC [37], a modern RML processing engine with a focus on speed and scalability, to execute the RML mapping rules and generate the RDF graph. Figure 4d shows an extract of the RDF graphs obtained by applying schema mapping rules to our example datasets. RDF graphs are staged in N-quads format, a serialization format for RDF data that associates each triple with an optional context value at the fourth position. This fourth field is used to track the provenance of the transformed data (identifier of the original data source).

### Data loading

The third and final step of the ETL data integration process involves saving the RDF graph generated during the data transformation stage into the triplestore. Different triplestore implementations may use different techniques to ingest RDF data. InteGraph provides connectors to the following triplestore solutions: RDFox, GraphDB, and Virtuoso. InteGraph also supports loading RDF data to a triplestore using SPARQL Update operations.

At the moment, inteGraph supports full load only. This means that the transformed data are loaded in full at each run of the ETL pipeline. Therefore, the KG is reconstructed from scratch every time the data integration pipelines are executed. A useful alternative would be incremental data load, i.e. updating the KG at regular intervals by loading only the data that has changed (new or updated data) since the last execution. This requires additional tools to compare the data from the data source with the existing data present in the KG. Incremental data load has a number of advantages over full load, including faster processing and preservation of data history.

### Pipeline creation, scheduling and monitoring

InteGraph uses Apache Airflow^2^ to create the data integration pipelines dynamically, and to schedule and monitor their execution. Airflow provides a flexible programmatic approach to easily build scheduled data processing pipelines as directed acyclic graphs (DAGs) of tasks. DAGs are a natural representation for ETL pipelines as each step in the ETL process is executed after the previous one has been completed (there is no circular dependency between ETL tasks). In addition, the extraction, transformation and loading steps can also be decomposed into sequences of lower-level atomic tasks. For instance, the transformation step involves tasks such as cleansing, taxonomic mapping, ontological mapping, and RDF graph generation. Therefore, each step of the ETL process can be implemented as a Airflow DAG, and the whole ETL pipeline as a DAG of DAGs.

Airflow supports parallel execution of independent tasks. Therefore, ETL pipelines can run in parallel, as each data source is independent from the other sources. In addition, independent tasks within a pipeline can also be parallelized. For example, inteGraph implements a mechanism to split an extracted data table into chunks that are transformed in parallel and merged before loading.

Airflow also enables to define a schedule interval for each pipeline, which determines when and how often the pipeline is run. In addition to scheduling and executing DAGs, Airflow provides a user interface to visualize the pipelines generated by inteGraph and monitor their execution (Figure 5). Airflow can also handle failures in ETL operations by retrying them a couple of times. If the error persists, the user can easily explore the logs of the failing task, identify the cause of the failure, and rerun the failing task (together with any subsequent tasks that depend on that task).

**Figure 5.**
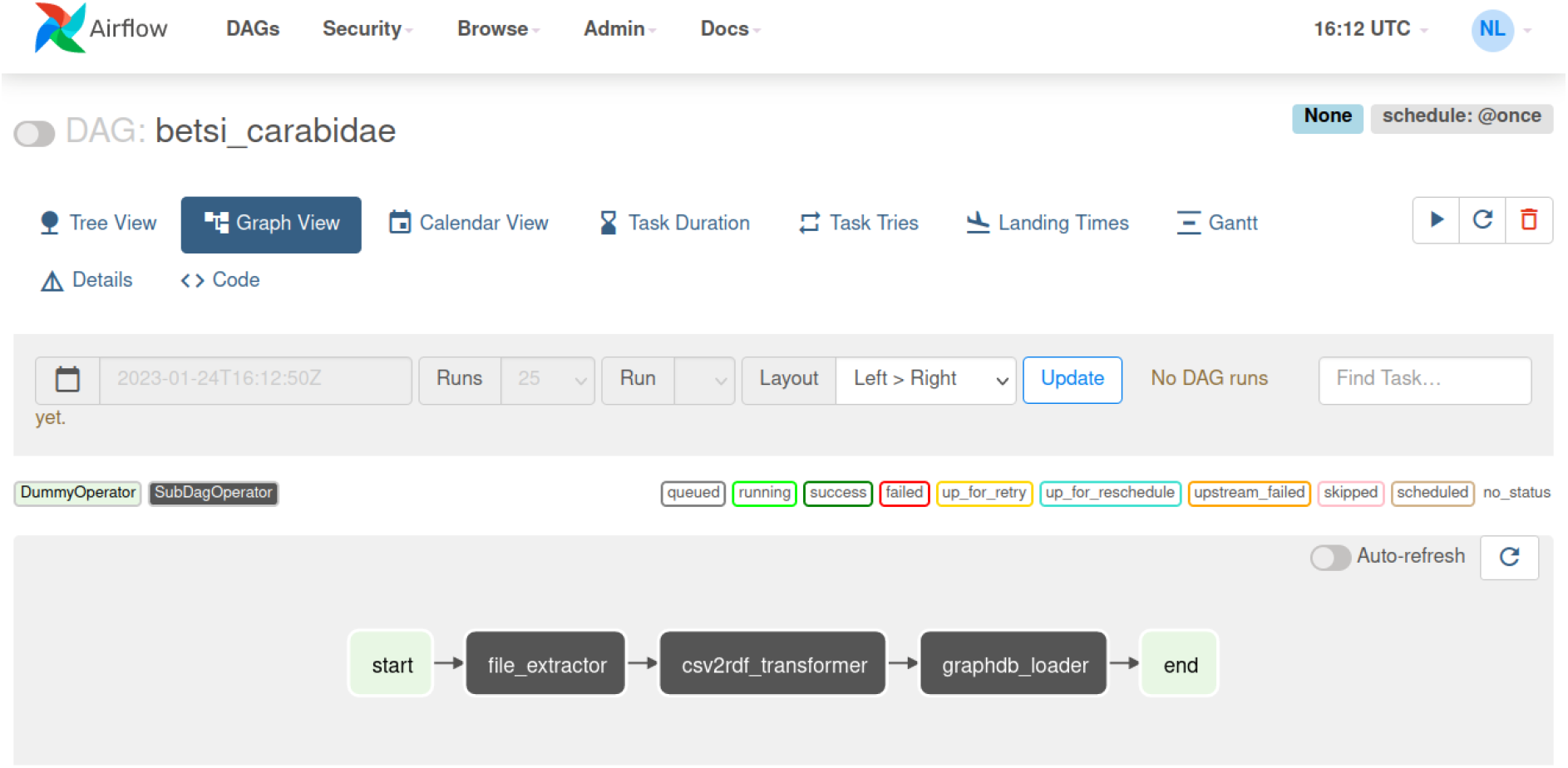
The ETL pipeline for the BETSI Carabidae dataset in Airflow user interface.

## Results

### Knowledge retrieval

At the end of the semantic data integration process, the target ontology and the transformed data are both saved in a single triplestore. The triplestore is responsible for storing the KG and executing SPARQL queries to retrieve information from it. SPARQL is a query language for retrieving and manipulating data stored in RDF format. SPARQL is based on matching graph patterns against the RDF graph. The basic graph pattern is the triple pattern, which is like a RDF triple where any part of the triple can be replaced by a variable. A graph pattern is a combination of such triple patterns. When executing a SPARQL query against a KG in a triplestore, the triplestore searches for the set(s) of triples that exactly match the graph patterns defined in the query, regardless of the provenance of the triples (unless explicitly requested). This means that the set of RDF triples returned in response to a SPARQL query may contain facts originating from different data sources. The KG provides the user with a unified view of the original data sources through querying, and enables combining multisource information as part of a query response (Figure 6).

**Figure 6.**
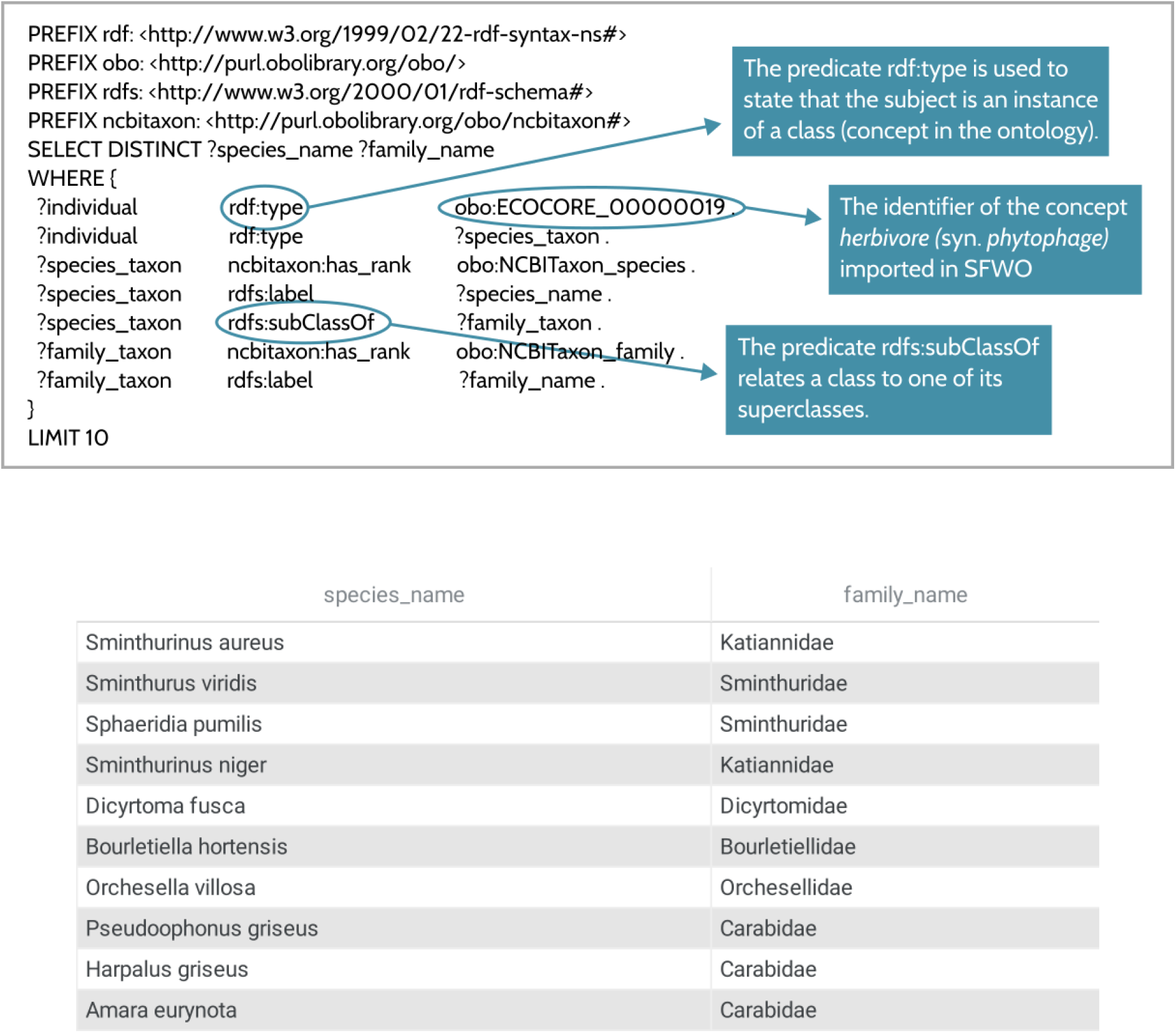
Example of a SPARQL query returning the species and family names of phytophagous taxa. ?x denotes a variable called x. The LIMIT keyword is used to limit the number of results to the first 10 entries. The query returns information about both phytophagous Collembola families (originating from the GloBi database) and phytophagous Carabidae (coming from the BETSI database).

### Making implicit knowledge explicit

The semantic data integration process builds a KG by linking heterogeneous datasets at the schema level using an ontology. During the process, the data layer of the KG is populated with the factual information stated in the different datasets and transformed into knowledge through semantic annotation and transformation into a RDF graph. Based on these explicit facts, additional knowledge that is not explicitly present in the data can be derived using reasoning. This ability is a direct consequence of OWL (the standard language for specifying ontologies) being based on a subset of first-order logic. Therefore, automated reasoners can be employed to evaluate the logical implications of the knowledge encoded in the ontology on the explicitly-stated data.

There are two principle strategies for logical inference: forward chaining and backward chaining [38]. Forward chaining, also known as materialization, materializes all the facts that can be logically deduced from the existing facts and a set of logical rules, and stores these inferred facts in the triplestore for later querying. Precomputing all inferred facts enables efficient query answering, but it can also be very expensive both in time (the materialization process needs to consider all possible inferences) and memory (the process can derive a large number of facts). In addition, materialization must be redone each time the data is updated.

Backward chaining (query rewriting) starts from a query and applies the logical rules only as far as they are needed to answer the query. With backward chaining, reasoning is done at runtime and no time- and space-consuming precomputation is needed. Furthermore, no recomputation has to be done when the data is updated. However, a major drawback of backward chaining is that reasoning must be done for each new query, which can be computationally expensive and slow.

In our running example, the target ontology and the transformed data are loaded in a triplestore supporting reasoning based on forward chaining. After data are loaded to the triplestore, the inference rules are applied repeatedly to the asserted (explicit) statements until no further inferred (implicit) statements are produced. Figure 7 illustrates how materialization makes implicit knowledge explicit in the KG. Given (1) the explicit information about *Entomobrya ligata* feeding on rotting wood (see first line of data table D3 in Figure 4a), (2) the hierarchy of taxonomic concepts provided by the NCBITaxon ontology, and (3) the logical definitions of diet concepts and trophic groups provided in the Soil Food Web Ontology, the triplestore reasoner is able to materialize the following logical implications:

- *E. ligata* is a species of springtails (Collembola);
- *E. ligata* is a detritivorous organism, as it feeds on rotten wood, which is a type of detritus;
- *E. ligata* belongs to the group of detritivorous springtails (Collembola.detritivores) as a logical consequence of the two previous assertions.

**Figure 7.**
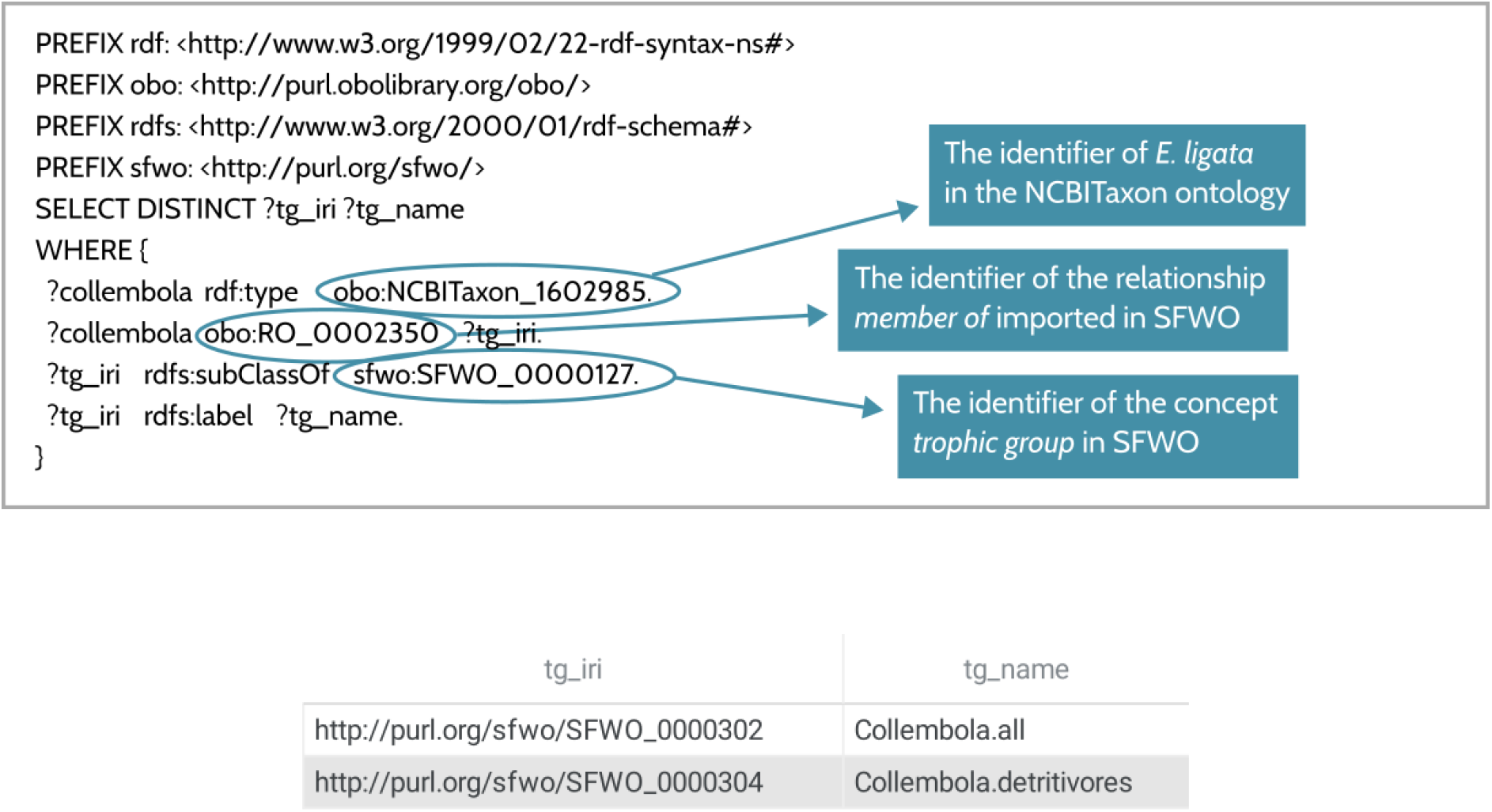
Example of a SPARQL query returning the trophic groups to which Entomobrya ligata belongs. A trophic group is defined in the Soil Food Web Ontology as “a collection of organisms that feed on the same food sources and have the same consumers”[7, 30]. SFWO provides a logical formalization of the hierarchical classification of soil consumers proposed in [39].

## Discussion

Multitrophic studies require integrating datasets across a large variety of taxonomic groups and trophic levels. Despite considerable efforts to make more biodiversity data freely available in a (semi-)structured format, the multiple dimensions of data heterogeneity (syntactic, structural, semantic) constitute a major obstacle to the interoperability of data sources [3]. Here, we introduced a practical approach to data integration that aims at making heterogeneous and distributed biodiversity data sources interoperable as part of a single KG. KGs provide a unified view over disparate data sources and allow for retrieving data across these sources using a single query. By using ontologies as global schemas, they add semantics to the integrated data, making it easier for humans and computers to interpret the data and for reasoners to infer additional facts. As seen from the examples discussed in the Results section, the ability to reason about the data in our trophic KG opens avenues for automatic classification of soil organisms, which can facilitate the reconstruction of consistent soil food webs from multisource data. In addition, KGs provide support for a number of applications [40], including both in-KG, e.g. link prediction, error detection, and out-of-KG applications, e.g. relation extraction from text, recommender systems, etc.

Despite their many advantages, KG construction is currently out-of-reach for most biodiversity data providers and consumers as they require in-depth expertise in Semantic Web technologies. InteGraph is an attempt to make semantic data integration and KG construction more accessible to the biodiversity science community. Requiring little or no code and minimal knowledge of the Semantic Web, inteGraph facilitates the processes of converting a biodiversity dataset into a KG and of integrating multiple datasets into a single KG. Given a set of distributed data sources and a target ontology, inteGraph allows to control the creation and execution of reproducible ontology-based data integration pipelines by providing a set of configuration files for each data source. This declarative approach relieves the user of the implementation burden. Instead, the user can focus on the schema of the input data, on the desired structure of the target KG, and on the schema mapping rules needed to transform the input data into RDF graphs. InteGraph relies on high-performance third-party tools (gnparser, nomer, Morph-KGC, Airflow), which guarantees a certain ability to scale to large datasets. The viability of our approach has been tested by creating a KG of soil trophic ecology from multiple open trait databases, using the Soil Food Web Ontology and the NCBITaxon ontology as the KG schema. The goal of this KG is to facilitate the realization of multitrophic studies in soil ecological research. Currently, inteGraph is at the proof-of-concept stage. It still needs some development to make it more robust, scalable and user-friendly. We also plan to add more advanced features in the future, especially regarding data provenance tracking and continuous KG updating.

Although it represents a significant advance in the field of ontology-based biodiversity data integration, inteGraph suffers limitations related to current practices in biodiversity data management. First, inteGraph requires the data sources to provide data in a (semi-)structured format through a programmatic interface, e.g. a URL to download the data file or a web API that handles HTTP requests. Still lots of data about soil biodiversity are not accessible this way, e.g. data from the BETSI database must be downloaded manually. We are confident that this situation will become less frequent in the future. Second, as a top-down approach to KG creation, inteGraph requires a predefined ontology to act as the mediating schema to link heterogeneous data sources. Creating an ontology to model knowledge in a domain of interest is a complex process that requires a significant investment of time and effort. Ontology engineering asks for a group of experts to produce a consensual conceptualization of the domain. For instance, in the domain of soil trophic ecology, this means trying to harmonize the use of diet terms that may have different meanings from one taxonomic group to another.

However, we believe that the result is worth the effort, as a properly designed ontology can benefit the whole community by facilitating knowledge sharing, dataset standardization and data integration.

Finally, although it aims to make semantic data integration more accessible to a non-expert audience, inteGraph still requires a minimum of knowledge of Semantic Web technologies (RDF data, ontologies, SPARQL queries…). Just as the environmental community has begun to embrace new artificial intelligence tools from recent developments in deep machine learning, we encourage the community to take an interest in Semantic Web tools for better biodiversity knowledge management.

Continuing our efforts to develop more and more biodiversity ontologies [41, 24, 42, 7] would allow us to envision increasing semantification of the ecology domain in the near future. Combined with tools such as inteGraph, which facilitate the conversion of biodiversity datasets into graph knowledge bases, these semantic resources could support the creation by different communities of numerous domain-specific KGs, which could eventually be interconnected to form a single biodiversity KG covering the entire tree of life and the full diversity of global ecosystems.

### Box 1. Glossary

**Extract-Transform-Load:** a three-phase data integration process that combines data from multiple sources into a single central repository.

**Knowledge graph:** a knowledge base that uses a graph-structured data model to integrate data.

**Ontology:** the formal and consensual description of a domain of interest as a set of interrelated concepts.

**Resource Description Framework (RDF):** the standard data model of the Semantic Web. RDF represents any piece of information as subject-predicate-object triples.

**RDF Mapping Language:** a language for expressing mapping rules from heterogeneous data structures to the RDF data model.

**Semantic data integration:** the process of combining data from different sources into a single, unified view using ontologies.

**Semantic Web:** a set of standard technologies - including the Resource Description Framework (RDF), Web Ontology Language (OWL), and SPARQL - that help make computers better able to interpret data and information published on the web.

**SPARQL:** the standard query language for retrieving and manipulating data stored in RDF format.

**Triplestore:** a database engine optimized for the storage and retrieval of RDF data.

**Web API:** an interface consisting of one or more endpoints publicly exposed on the web, that allow a user to programmatically access some specific features or the data of an application, e.g. a database.

**Web Ontology Language (OWL):** a family of knowledge representation languages for authoring ontologies, built on RDF and characterized by formal semantics based on description logics (decidable fragments of first-order logic).

## Acknowledgements

We acknowledge support from the European Union’s Horizon Europe under grant agreement N°101060429 (NaturaConnect), the French Agence Nationale de la Recherche through the EcoNet (ANR-18-CE02-0010), GlobNet (ANR-16-CE02-0009) and FishPredict projects, the MIAI@Grenoble Alpes (ANR-19-P3IA-0003) institute, and the Investissement d’Avenir grants (Trajectories: ANR-15-IDEX-02; Montane: OSUG@2020: ANR-10-LAB-56) and the METRO Grenoble Alpes and the Isere department.

## Footnotes

1 Available at https://github.com/nleguillarme/inteGraph

2 https://airflow.apache.org/

